# Evolution of bird communication signals: transference between signals mediated by sensory drive

**DOI:** 10.1101/142463

**Authors:** Oscar Laverde-R., Michael J. Ryan, Carlos Daniel Cadena

## Abstract

Animals communicate using signals perceived via multiple sensory modalities but usually invest more heavily in one of type of signal. This pattern, observed by Darwin^1^ and many researchers since, led to development of the transfer hypothesis (see also transferal effect^2^ and tradeoff hypothesis^3,4^), which predicts a negative relationship between investment in different signaling modalities dictated by the relative costs and benefits of each. One factor that influences costs and benefits, and is central to the sensory drive hypothesis^5^ posed to account for signal evolution, is the suitability of the environment for different types of signals. Movement into a dark habitat, for example, should favor investment in acoustic over visual signals. We use phylogenetic comparative methods to analyze the joint effect of transfer and sensory drive on plumage and song variation in 52 species of a large radiation of passerine birds, the New World warblers (Parulidae), and to estimate temporal patterns in the accumulation of differences in visual and vocal signals and habitat along the evolutionary history of this lineage. We found evidence for the predicted negative correlations between a variety of song and plumage traits that vary with habitat type. Plumage contrast to background and chromatic diversity were both negatively related to syllable variety when vegetation structure was a covariate: birds with a greater variety of song syllables and less colorful plumages live in closed or darker habitats. Also as predicted, achromatic or brightness diversity was related to vegetation structure. In addition, disparity-through-time analyses showed that when one set of traits (i.e. songs or colors) diversified at a relatively high rate the other did not, as predicted by the transfer hypothesis. Our results show that sensory drive influences the transfer of investment between traits in different sensory modalities. This interaction between mechanisms shaping signals may be a major determinant in the evolution of animal communication.

---

Ever since Darwin^1^, naturalists have noted a negative relationship between bright colors and the elaboration of songs involved in animal communication. For example, “…*in territorial passerine birds, there tends to be an inverse relation between the development of auditory distinctiveness (song of species with cryptic behavior and coloration, e.g., grasshopper warbler, chiffchaff) and visual distinctiveness (pattern of species with conspicuous behavior, e.g., stonechat, pied flycatcher)*…” ^6^. Such observations led to the postulation of the transferal effect^2,7^ or tradeoff hypothesis^3^ (later named transfer hypothesis), which states that the complexity of signals in different modalities should be negatively related. Animals are expected to invest primarily in one type of signal (i.e., vocal, chemical, or visual^8^) owing to various factors, including internal limitations in energy expenditure (classical life-history tradeoffs^9,10^), predators^11^, parasites^12^, physical conditions of the habitat^2^, and the conflicting forces of sexual and natural selection^12^. Despite the generality and popularity of this hypothesis, there have been few^13-15^ attempts to test it with a robust, phylogenetically informed data set that includes variation in multiple signal modalities and covariation with the signaling habitat that influences the salience of these signals.

Here, we test for a negative relationship between elaboration in song and plumage coloration in the context of the signaling habitat, thus linking the transfer hypothesis with another leading hypothesis posed to account for signal evolution, namely sensory drive^5^. The sensory drive hypothesis posits that variation in habitat features leads to variation in selection pressures by affecting the ease with which different traits are perceived. For example, because the use of visual signals for communication in dark habitats (e.g. forest understory, turbid waters) is expected to be ineffective^5,16^, acoustic or olfactory signals should be more prevalent as targets of sexual selection in such environments. In turn, in open areas or clear waters one expects visual signals to be more elaborate than vocal or chemical signals. Although several studies have supported the sensory drive hypothesis, they have largely focused on a single communication channel (i.e., visual^15,17-20^, olfactory^16^, or acoustic signals^21-23^) and have not considered interactions among channels as those implied by the transfer hypothesis. Similarly, the few existing examinations of the transfer hypothesis^13-15,24^ have not considered the influence of habitat on signal salience.

Using data on plumage, song, and habitat, we examined correlations between visual and acoustic signals mediated by the environment across a large radiation of passerine birds. The New World warblers (Parulidae) are a group of small, often colorful, primarily insectivorous oscines with diverse songs and a broad diversity of habitat affinities and life histories^25^. The tuning of the visual system of warblers varies with habitat and this should generate differential, habitat-specific selection on visual signals. Expression of opsins varies with the light environment occupied by different species^26^; specifically, short-wavelength sensitive type 2 opsins (SWS2) are shifted to longer wavelengths in species living in closed forest environments^27^ where most of the short wavelengths are filtered by vegetation^28^. In addition, female expression of these same opsins varies with plumage dichromatism, suggesting a role for sensory drive^26^.

In analyses that did not account for the habitat preferences of species, we found no support for the prediction of the transfer hypothesis that elaboration in plumage coloration (chromatic diversity of plumage and chromatic contrast to the back) and song traits (vocal deviation -a measure of vocal performance-, syllable variety and song length) should be negatively related (Table 1, 2). However, when we included vegetation structure as a covariate (i.e., open vs. closed habitats), significant negative relationships between variables emerged, suggesting that apparent transfers in investment between signals in different communication channels are mediated by habitat. Specifically, chromatic contrast to background (β= -6.79, t= -2.69, p=0.01) and chromatic diversity of plumage were negatively related to song syllable variety (β= -1.51, t= -3.31, p=0.001, Table 1), such that species with less colorful plumage exhibit songs with a greater variety of syllables and live in closed habitats (β= 1.96, t= 2.76, p=0.008). Additionally, achromatic diversity of plumage was significantly related to vegetation structure (β= 3.22, t= -2.16, p=0.03, Table 2) and to vocal deviation (β=1.6290, t=2.0145, p=0.05, Fig. 1); this indicates that birds with darker backs live in darker habitats and communicate using songs demanding greater performance. A plausible interpretation of these results is that in forest species natural selection to reduce conspicuousness due to predation has resulted in a match between plumage coloration and background color; this has presumably triggered an investment transfer between visual and vocal signals, resulting in more elaborate songs favored by sexual selection. These patterns we found are uniquely predicted by a combination of the transfer and sensory drive hypotheses.

**Figure 1.**
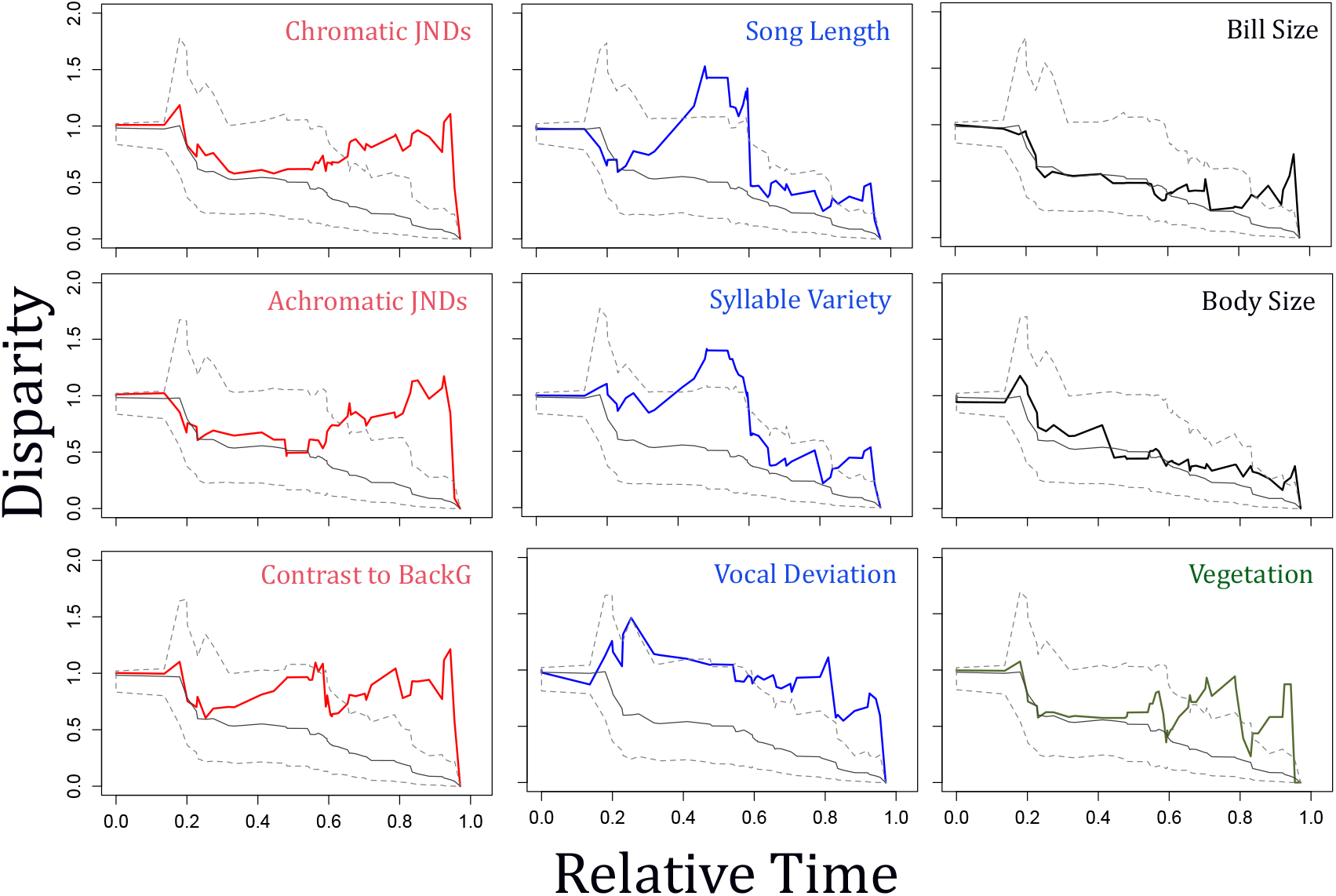
Disparity through time (DTT) plots for plumage, songs, morphology and vegetation suggest a negative evolutionary correlation between plumage traits and attributes of songs possibly mediated by habitat. Disparity in color traits (in red) and habitat (in green) accumulated at a faster rate near the tips of the phylogeny.However, disparity in variables related to song elaboration (in blue) peaked around the middle of the clade’s history. Morphological traits like body size and bill size (in black) followed a null evolutionary model.

**Table 1.** Phylogenetic generalized least squares multiple regressions of plumage contrast to the back on different sets of predictors. Only when we included vegetation structure as a covariate did significant negative relationships between variables emerge (the model that best fit the data is shown in bold), suggesting that transfers in investment between signals in different communication channels are mediated by habitat.

| Independent variables | AIC | p-value | Adjusted R-squared |
| --- | --- | --- | --- |
| Syllable variety | 84.18 | 0.89 | -0.02 |
| Vocal Deviation | 84.28 | 0.98 | -0.02 |
| Song Length | 83.90 | 0.68 | -0.01 |
| Vegetation | 83.95 | 0.71 | -0.02 |
| **Syllable Variety \* Vegetation** | **76.77** | **0.01** | **0.17** |
| Song Length * Vegetation | 87.36 | 0.88 | -0.05 |
| Vocal Deviation * Vegetation | 86.95 | 0.80 | -0.04 |

**Table 2.** Phylogenetic generalized least squares multiple regressions of achromatic diversity (dL) on different sets of predictors. Achromatic diversity of plumage was significantly related to vocal deviation and vegetation structure. Vocal deviation was a good predictor of achromatic diversity, but the explanatory power of the model increased when vegetation was included in the model. This indicates that birds with darker backs live in darker habitats and communicate using songs demanding greater performance.

| Independent variables | AIC | p-value | Adjusted R-squared |
| --- | --- | --- | --- |
| Syllable variety | 283.99 | 0.32 | 0.00 |
| **Vocal Deviation** | **281.67** | **0.02** | **0.07** |
| Song Length | 284.30 | 0.44 | 0.00 |
| **Vegetation** | **280.76** | **0.02** | **0.10** |
| Syllable Variety * Vegetation | 283.52 | 0.14 | 0.12 |
| Song Length * Vegetation | 282.39 | 0.08 | 0.08 |
| **Vocal Deviation \* Vegetation** | **281.69** | **0.03** | **0.12** |

A more specific prediction of the transfer hypothesis considers not only the evolutionary end points of elaboration in visual versus vocal signals, but the dynamic processes that gave rise to their negative relationship. Analyses of the accumulation of disparity in color and vocal traits along the New World warbler phylogeny suggest a negative evolutionary correlation between these two different types of signals across time. Relative to a null process of accumulation of phenotypic disparity (i.e., under a Brownian-motion process of evolution), chromatic (chromatic diversity of plumage and chromatic contrast to background) and achromatic (achromatic diversity of plumage) disparity in coloration accumulated at a faster rate near the tips of the phylogeny (Fig. 1), indicating divergence among closely related taxa. Accumulation in disparity in song length and syllable variety also differed from the null but, by contrast, peaked around the middle of the clade’s history and not near the present. Taken together, patterns of evolutionary change in plumage and song traits appear to be decoupled: when there were bursts in accumulation in disparity in song length and syllable variety, disparity in plumage coloration did not deviate from the null and *vice versa* (Fig. 1). In other words, it appears that when one set of traits diversified at a relatively high rate the other did not, as predicted by the transfer hypothesis. The pattern of disparity through time estimated for vegetation structure closely mirrored patterns observed for plumage coloration variables (Fig. 1). This suggests that diversification in coloration in warblers may have been linked to shifts between habitats differing in light availability and color properties as predicted by the sensory drive hypothesis. In contrast to traits involved in visual and acoustic communication, morphological traits (i.e., body mass and bill size) accumulated disparity as expected under a Brownian-motion model of evolution.

Considering vegetation structure allowed us to reveal a negative association between vocal and visual signals that was otherwise absent when vegetation structure was not considered a covariate. We thus conclude that the signaling environment mediates the evolution of the relationship between vocal and visual signals, resulting in negative relationships between the degree of elaboration of different types of signals (i.e., apparent transferal effects). A variety of studies have tested the transfer hypothesis in birds, with mixed results: finches and avenue-building bowerbirds exhibit negative relationships between acoustic and visual signals^24,29^, but no evidence of such apparent transference has been found in trogons^15^, Asian barbets^14^ or tanagers^13^. In other groups like butterflies^30^, fishes^16,31^ and plants^32^, there is evidence of negative relationships between signals driven by the properties of the habitat. For example, in turbid waters sticklebacks rely more on olfactory cues to attract mates than in clear waters^31,33^, suggesting a transference from visual signals to chemical signals, mediated by habitat.

Our study offers strong support for the transfer hypothesis, but only when it is considered in the context of sensory drive. Transfer in signal investment occurs when there are differential costs and benefits of signals in different modalities. Our results suggest that these differential costs and benefits need not occur because of internal physiological or developmental constraints as previously suggested, but can be generated by the saliency of signals in their local habitat. We conclude that to understand the diversity of multimodal communication strategies it is necessary to address not only the magnitude of investment in signaling modalities but also the selective forces that influence their salience.

## METHODS SUMMARY

We measured coloration in New World warblers using museum specimens with an Ocean Optics (Dunedin, FL) USB2000 spectrophotometer and a PX-2 pulsed xenon light source to record reflectance across the avian visual spectrum. For each specimen, we measured all plumage patches we observed that appeared to be differently colored to the human eye along 40 measurement points in ventral and dorsal sides. With these data, we constructed three variables describing colors: (1) chromatic and (2) achromatic diversity, and (3) contrast to background, assuming that the color in of the backs matched the color of the background, we can use that variable as a proxy of the color of the background where birds live. Chromatic and achromatic diversity were calculated by comparing the contrast between every color measured on each specimen using the library pavo^47^, implemented in R.

We used data on song length, syllable variety and vocal deviation from a previous study^46^ as measurements of song elaboration^13^ in New World warblers. We explored the evolutionary history of communication signals (visual and acoustic) in New World warblers (Parulidae) using comparative methods designed to understand the patterns of evolution of traits along a phylogeny^34^ (DTT plots) and phylogenetic generalized least squares (PGLS)^35^ models to test for relationships between variables predicted by the transfer hypothesis and sensory drive.

## METHODS

We explored the evolutionary history of communication signals using comparative methods designed to understand the patterns of evolution of traits along a phylogeny^34^ (disparity through time [DTT] plots) and phylogenetic generalized least squares (PGLS)^35^ to evaluate different models. We focused on New World warblers (Parulidae), which have evolved in contrasting habitat types, and differ in communication strategies to study the evolution of communication signals.

### Plumage conspicuousness

We measured coloration in New World warblers using museum specimens. In total we measured 231 study skins of 51 species (see Supplement) in the collections at the Museum of Natural Science at Louisiana State University (LSUMZ). To avoid any effects of specimen age on reflectance measures^17,36^ the most recent specimens with the freshest looking plumage for each species were chosen. We used an Ocean Optics (Dunedin, FL) USB2000 spectrophotometer with a PX-2 pulsed xenon light source to record reflectance across the avian visual spectrum. All measurements were taken at a 45-degree angle to the feather surface. For each species and each sex, we measured all colors we observed in all patches that appeared to be differently colored to the human eye along on 40 measurement points in the ventral and dorsal sides of each specimen. These measurements encompassed all major plumage patches (dorsal: crown, mantle and rump; ventral: throat, breast and belly).

To quantify the conspicuousness of plumage coloration we used two approaches. First, we calculated euclidean distances between the colors in the belly, rump and crown against the color in the mantle. Birds increase crypsis by displaying countershaded patterns (lighter ventral than dorsal parts) and by matching background contrast (Fig. S1), particularly on their dorsal parts, which are often more exposed to predators ^28,37^. For this reason, being unable to measure light environments for all species in the field, we assumed that dorsal plumage approximately emulates –or at least more closely resembles– the color of the background where birds live.

Therefore, the contrast between the back and the belly, crown and rump colors was used as a proxy for conspicuousness. For this approach, we used an avian visual model^38,39^ to analyze colors in a tetrahedral color space. We called this variable contrast to the back. Our second approach used a discrimination model which calculates a contrast, for the chromatic and achromatic domain, in avian color space defined by the quantum catches of each receptor type (i.e., cone cell) in the avian retina^40^. The model assumes that color discrimination in this perceptual space is limited by noise originating in the receptors, and that no visual signal results when stimulus and background differ only in intensity^40^. We calculated the chromatic (dS) and achromatic (dL) diversity of plumages as follows. First, we measured the contrasts between every pair of points (40) measured on each specimen; second, we calculated the centroid for the chromatic and the achromatic dimension; third, we measured the distance from each point to the centroid for the chromatic and achromatic measures of contrast and we averaged these measurements to have a mean value of the distribution of the contrasts in the chromatic and achromatic dimension of colors (Fig S2). We called these measurements achromatic (dL) and chromatic (dS) diversity of plumage.

### Song measurements

We used data on song length, syllable variety and vocal performance from a previous study^41^ as measurements of song elaboration in New World warblers^13^. We also used existing data on bill size^41^ and body size^42^ to study the evolution of morphological traits, compared to the evolution of communication traits.

As songs increase in either frequency bandwidth or trill rate, the demands of vocal performance increase. Vocal deviation (i.e., the orthogonal distance from the upper - bound regression line between frequency bandwidth and pace of each trill) is strongly related to the energetic demands of vocal performance^43,44^. Songs far below the regression line (i.e. those with higher values of vocal deviation) are assumed to be less vocally demanding than those closer to the regression line^44^; thus, we interpret lower values of vocal deviation to reflect greater investment in song production.

### Vegetation structure

Vegetation structure was scored using habitat descriptions^45^ and in accordance with previous work^41^, as follows: 1 – open, 2 – semiclosed with low vegetation, 3 – semiclosed with high vegetation and 4 – closed. Intermediate scores were used when species used more than one habitat type.

### Analyses

To evaluate the transfer hypothesis and the effect of the vegetation density on how transferences are resolved, we used a generalized least square model for comparative phylogenetics (PGLS) implemented in the package caper^46^. We run PGLS setting contrast to background, chromatic (dS) and achromatic (dL) diversity of plumage as response variables including different set of predictors. First, to test for transfer effects we run models independently with syllable variety, vocal deviation and song length as predictors. Then, to test the effect of habitat on the relationship between visual and acoustical signals, we added vegetation structure as another predictor in the models.

To examine patterns of evolutionary differentiation in vocal performance and color conspicuousness among lineages of New World warblers, we constructed disparity-through-time (DTT) plots^34^. DTT plots allow one to examine the accumulation in trait disparity over evolutionary time in a clade by estimating the dispersion of points in multivariate space across time intervals in a phylogeny. This method calculates dissimilarity as the mean square (pair-wise) Manhattan distance among points in trait-space for subclades relative to the variance of the entire clade. Disparity is calculated for individual nodes by moving up the phylogeny from the root of the tree. Relative disparity values close to 0.0 indicate that subclades contain only a small proportion of the total variation and therefore overlap in occupation of phenotypic space is minimal between the different subclades; conversely, relative disparity values close to 1.0 indicate extensive phenotypic overlap^34^. For all the comparative analyses mentioned above, we used a molecular phylogeny of New World warblers based on several mitochondrial and nuclear gene sequences^25^.

---

**Figure S1.**
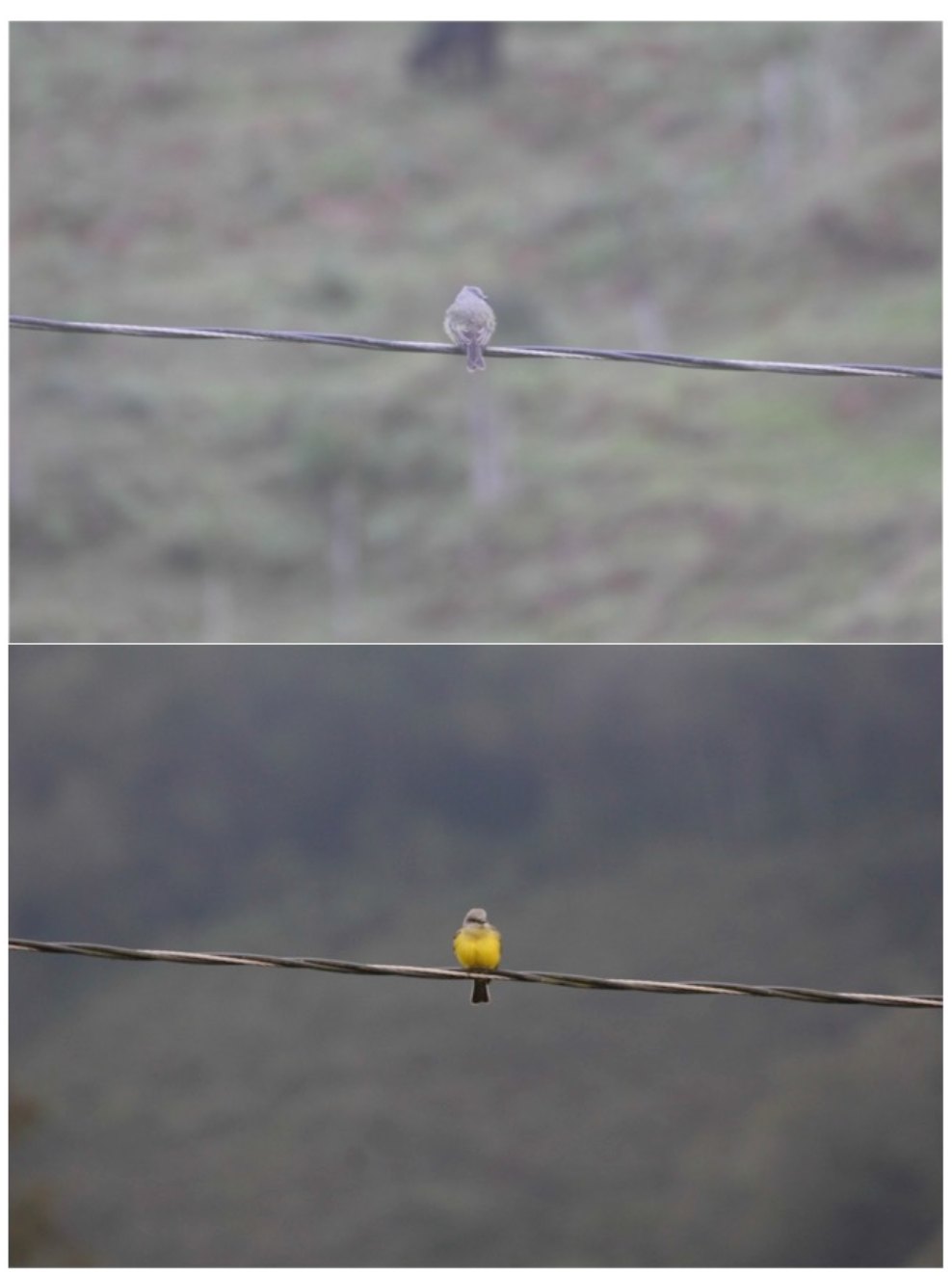
Two photographs of the Tropical Kingbird *(Tyrannus melancholichus)* as an example of how dorsal colors match the background color while the ventral color is conspicuous relative to the background. We assume background color matching by the dorsal parts as shown in this bird also occurs in the New World warblers.

**Figure S2.**
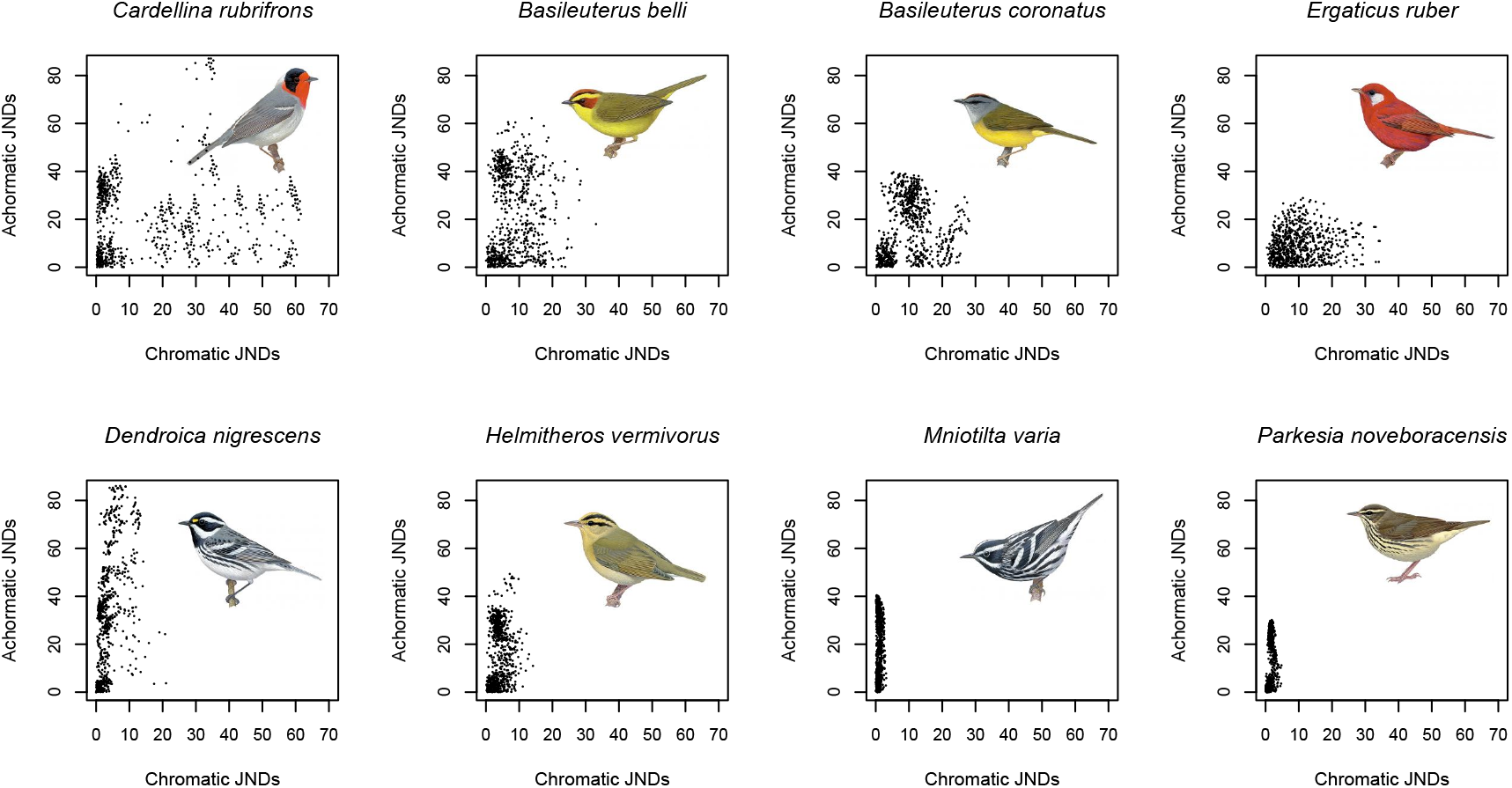
Variation in achromatic and chromatic diversity of colors among selected species of warblers. Each point is a comparison between the discrimination thresholds or ‘just noticeable differences’ (JND) between two reflectance measures. Values in the 1–3 range indicate that two objects are likely to be indistinguishable to an avian observer, whereas values >3 are increasingly likely to lead to detection and discrimination. Notice the distribution of points in *Mniotilta varia* and *Parkesia noveboracensis,* which exhibit wide variation in the achromatic axis relative to low variation in the chromatic axis. In contrast, *Cardellina rubrifrons* and *Basileuterus belli* exhibit ample variation along both achromatic and chromatic axes.

## Literature cited

1. Darwin, C. The Descent of Man and Selection in Relation to Sex. London: John Murray. (1871).

2. Gilliard, T. Bower ornamentation versus plumage characters in Bowerbirds. Auk 73, 450–451 (1956).

3. Repentigny, Y. D. E., Ouellet, H. & McNeil, R. Song versus plumage in some North American oscines: testing Darwin’s hypothesis. Ecoscience 7, 137–148 (2000).

4. Shutler, D. Sexual selection: when to expect trade-offs. Biol. Lett. 7, 101–104 (2011).

5. Endler, J. A. Signals, signal conditions, and the direction of evolution. Am. Nat. 139, S125–S153 (1992).

6. Huxley, J. S. Darwin’s theory of sexual selection and the data subsumed by it, in the light of recent research. Am. Nat. 72, 416–433 (1938).

7. Gilliard, T. E. The evolution of bowerbirds evolutionary processes. Sci. Am. 209, 38–46 (1963).

8. Gil, D. & Gahr, M. The honesty of bird song: multiple constraints for multiple traits. Trends Ecol. Evol. 17, 133–141 (2002).

9. Zera, A. J. & Harshman, L. G. The physiology of life history trade-offs in animals. Annu. Rev. Ecol. Syst. 32, 95–126 (2001).

10. Roff, D. a & Fairbairn, D. J. The evolution of trade-offs: where are we? J. Evol. Biol. 20, 433–447 (2007).

11. Tuttle, M. D. & Ryan, M. J. Bat predation and the evolution of frog vocalizations in the Neotropics. Science 214, 677–678 (1981).

12. Partan, S. R. & Marler, P. Issues in the classification of multimodal signals. Am. Nat. 166, 231–245 (2005).

13. Mason, N. A., Shultz, A. J. & Burns, K. J. Elaborate visual and acoustic signals evolve independently in a large, phenotypically diverse radiation of songbirds. Proc. R. Soc. B Biol. Sci. 281, 1–10 (2014).

14. Gonzalez-Voyer, A., Den Tex, R.-J., Castelló, A. & Leonard, J. A. Evolution of acoustic and visual signals in Asian barbets. J. Evol. Biol. 26, 647–659 (2013).

15. Ornelas, J. F., González, C. & Espinosa de los Monteros, A. Uncorrelated evolution between vocal and plumage coloration traits in the trogons: a comparative study. J. Evol. Biol. 22, 471–484 (2009).

16. Rafferty, N. E. & Boughman, J. W. Olfactory mate recognition in a sympatric species pair of three-spined sticklebacks. Behav. Ecol. 17, 965–970 (2006).

17. Doucet, S. M., Mennill, D. J. & Hill, G. E. The evolution of signal design in manakin plumage ornaments. Am. Nat. 169, S62–S80 (2007).

18. Cardoso, G. C. & Mota, P. G. Evolution of female carotenoid coloration by sexual constraint in Carduelis finches. BMC Evol. Biol. 10, 82 (2010).

19. Price, J. J. & Whalen, L. M. Plumage evolution in the oropendolas and caciques: different divergence rates in polygynous and monogamous taxa. Evolution 63, 2985–2998 (2009).

20. Mcnaught, M. K. & Owens, I. Interspecific variation in plumage colour among birds: species recognition or light environment? J. Evol. Biol. 15, 505–514 (2002).

21. Seddon, N. Ecological adaptation and species recognition drives vocal evolution in neotropical suboscine birds. Evolution 59, 200–215 (2005).

22. Kirschel, A. N. G. et al. Birdsong tuned to the environment: green hylia song varies with elevation, tree cover, and noise. Behav. Ecol. 20, 1089–1095 (2009).

23. Slabbekoorn, H. & Smith, T. B. Habitat-dependent song divergence in the little greenbul: an analysis of environmental selection pressures on acoustic signals. Evolution 56, 1849–1858 (2002).

24. Badyaev, A. V, Hill, G. E. & Weckwort, B. V. Species divergence in sexually selected traits: increase in song elaboration is related to decrease in plumage ornamentation in finches. Evolution 56, 412–419 (2002).

25. Lovette, I. J. et al. A comprehensive multilocus phylogeny for the wood-warblers and a revised classification of the Parulidae (Aves). Mol. Phylogenet. Evol. 57, 753–770 (2010).

26. Bloch, N. I. Evolution of opsin expression in birds driven by sexual selection and habitat. Proc. R. Soc. B Biol. Sci. 282, 20142321, (2014).

27. Bloch, N. I., Morrow, J. M., Chang, B. S. W. & Price, T. D. SWS2 visual pigment evolution as a test of historically contingent patterns of plumage color evolution in warblers. Evolution 69, 341–356 (2015).

28. Gomez, D. & Théry, M. Simultaneous crypsis and conspicuousness in color patterns: comparative analysis of a Neotropical rainforest bird community. Am. Nat. 169, S42–S41 (2007).

29. Kusmierski, R., Borgia, G., Uy, A. C. & Crozier, R. H. Labile evolution of display traits in bowerbirds indicates reduced effects of phylogenetic constraint. Proc. Biol. Sci. 264, 307–313 (1997).

30. Vane-Wright, R. I. & Boppre, M. Visual and chemical signalling in butterflies: functional and phylogenetic perspectives. Philos. Trans. R. Soc. B Biol. Sci. 340, 197–205 (1993).

31. Heuschele, J., Mannerla, M., Gienapp, P. & Candolin, U. Environment-dependent use of mate choice cues in sticklebacks. Behav. Ecol. 20, 1223–1227 (2009).

32. Raguso, R. A. Wake up and smell the roses: the ecology and evolution of floral scent. Annu. Rev. Ecol. Evol. Syst. 39, 549–569 (2008).

33. Heuschele, J. & Candolin, U. An increase in pH boosts olfactory communication in sticklebacks. Biol. Lett. 3, 411–413 (2007).

34. Harmon, L. J., Schulte, J. a, Larson, A. & Losos, J. B. Tempo and mode of evolutionary radiation in iguanian lizards. Science 301, 961–964 (2003).

35. Freckleton, R. P., Harvey, P. H. & Pagel, M. Phylogenetic analysis and comparative data: a test and review of evidence. Am. Nat. 160, 712–726 (2002).

36. Doucet, S. M. & Hill, G. E. Do museum specimens accurately represent wild birds? A case study of carotenoid, melanin, and structural colours in long-tailed manakins Chiroxiphia linearis. J. Avian Biol. 40, 146–156 (2009).

37. Clarke, J. M. & Schluter, D. Colour plasticity and background matching in a threespine stickleback species pair. Biol. J. Linn. Soc. 102, 902–914 (2011).

38. Goldsmith, T. H. Optimization, constraint and history in the evolution of eyes. Quaterly Rev. Biol. 65, 281–322 (1990).

39. Endler, J. A. & Mielke, P. W. Comparing entire colour patterns as birds see them. Biol. J. Linn. Soc. 86, 405–431 (2005).

40. Vorobyev, M. & Osorio, D. Receptor noise as a determinant of colour thresholds. Proc. R. Soc. B Biol. Sci. 265, 351–358 (1998).

41. Cardoso, G. C. & Hu, Y. Birdsong performance and the evolution of simple (rather than elaborate) sexual signals. Am. Nat. 178, 679–686 (2011).

42. Dunning, J. B. J. CRC Handbook of Avian Body Masses. (CRC Press, 2008).

43. Ballentine, B. Vocal performance influences female response to male bird song: an experimental test. Behav. Ecol. 15, 163–168 (2004).

44. Podos, J. A Performance constraint on the evolution of trilled vocalizations in a songbird family (Passeriformes: Emberizidae). Evolution 51, 537–551 (1997).

45. Curson, J., Quinn, D. & Beadle, D. The Warblers of the Americas. An Identification Guide. Houghton Mifflin, 1994.

46. Orme, D. et al. Caper: comparative analyses of phylogenetics and evolution in R. (2012).

47. Maia, R., Eliason, C. M., Bitton, P.-P., Doucet, S. M. & Shawkey, M. D. Pavo: an R package for the analysis, visualization and organization of spectral data. Methods Ecol. Evol. 906–913 (2013).

## References

36. Doucet, S. M. & Hill, G. E. Do museum specimens accurately represent wild birds? A case study of carotenoid, melanin, and structural colours in long-tailed manakins *Chiroxiphia linearis*. J. Avian Biol. 40, 146–156 (2009).

39. Endler, J. A., Westcott, D. A., Madden, J. R. & Robson, T. Animal visual systems and the evolution of color patterns: sensory processing illuminates signal evolution. Evolution 59, 1795–1818 (2005).

42. Dunning, J. B. J. CRC Handbook of Avian Body Masses. (CRC Press, 2008).

45. Curson, J., Quinn, D. & Beadle, D. The Warblers of the Americas. An Identification Guide. (Houghton Mifflin, 1994).

46. Orme, D. et al. Caper: Comparative analyses of phylogenetics and evolution in R. (2012).

